# GEMS (GWAS Effect Mechanisms via Sequence-to-Function): Unveiling cell type-specific mechanisms for GWAS variants

**DOI:** 10.64898/2026.10.14.759213

**Authors:** Xiaoman Xie, Sheng Qian, Diana Chang, Tushar Bhangale, Mark I. McCarthy, Gokcen Eraslan, Kipper Fletez-Brant

## Abstract

Genome-wide association studies (GWAS) have linked thousands of loci to common diseases, yet for non-coding variants, which constitute the vast majority of the signal, the cell types and states through which individual risk variants act remain largely unknown. Here, we develop GEMS (GWAS Effect Mechanisms via Sequence-to-Function), a framework that trains supervised classifiers on multimodal variant effect predictions (ATAC-seq and RNA-seq) from the sequence-to-function model Borzoi and applies SHAP-based attribution to associate individual GWAS variants with the cell types most relevant to their disease association. Applied to asthma, LDL cholesterol, multiple sclerosis, and type 2 diabetes, GEMS prioritizes known disease-relevant cell types concordant with stratified LD-score regression. At the variant level, GEMS associates a median of 58.7% of fine-mapped variants with at least one cell type via ATAC-seq and 65.9% via RNA-seq, extending cellular annotation beyond the chromatin accessibility peaks in the single-cell atlas used for model training. ATAC-seq variant–cell-type associations are significantly enriched in cis-regulatory elements from independent experimental studies not used in model training, and enrichment persists after excluding variants overlapping training atlas peaks. For multiple sclerosis, hierarchical biclustering of variant–cell-type associations partitions risk variants into immune and CNS modules driven by distinct transcription factor families, recapitulating the immune-mediated and neurodegenerative components of the disease and demonstrating that integrative analysis of sequence-to-function predictions can stratify phenotype-associated variants into mechanistically distinct, cell-type-specific regulatory programs.

## INTRODUCTION

Large-scale genome-wide association studies (GWAS) have now catalogued thousands of risk loci across common diseases and complex traits^1,2^, yet for most loci the cell types through which risk variants exert their effects remain unknown. The vast majority of GWAS associations map to non-coding regions of the genome^3,4^, where they are presumed to act through cell-type-specific modulation of gene expression^3,5^. Resolving which variants act through which cell types can stratify disease-risk loci into mechanistically distinct subgroups, distinguish individuals by the cellular contexts disrupted by their risk alleles, identify effector genes, and prioritize causal variants within unresolved credible sets.

Existing strategies for connecting GWAS variants to cellular context fall into two categories. At the variant level, colocalization with cell-type-specific expression quantitative trait loci (eQTLs) can associate variants with their relevant tissue or cell types and nominate effector genes at individual loci^6,7^. However, eQTL studies and GWAS capture fundamentally different components of the genetic architecture of complex traits^8^, and most GWAS loci do not colocalize with an eQTL in any tested tissue^7,9,10^. Annotating GWAS variants by overlap with single-cell epigenomic maps^11,12^ provides cell-type specificity but is constrained by incomplete cell-type representation and sparse peak coverage in single-cell chromatin profiles^13^, leaving many GWAS variants outside detected regulatory peaks^14^. At the phenotype level, stratified LD-score regression (sLDSC)^15^ tests whether disease heritability is enriched in regulatory elements active in specific cell types, and single-cell disease relevance scoring (scDRS)^16^ evaluates whether individual cells exhibit excess expression of GWAS-associated genes. These approaches have identified disease-relevant cell types across a range of phenotypes, but they quantify enrichment at the level of the trait as a whole and cannot resolve per-variant cellular contexts.

Sequence-to-function (S2F) deep learning models, including Enformer^17^, Borzoi^18^, and AlphaGenome^19^, offer a potential path forward. These models are trained to predict molecular phenotypes from DNA sequence alone, thereby learning a compressed representation of cell-type-specific regulatory logic that can be queried for variant effect predictions (VEPs) across thousands of cellular contexts simultaneously. Because they capture regulatory patterns on the sequence level, they bypass experimental data sparsity to offer a genome-wide assessment of the regulatory potential of genetic variants. However, raw S2F predictions represent a rich, high-dimensional feature space that requires systematic meta-modeling to disentangle trait-specific cellular context from background regulatory overlap^20,21^.

Here, we developed GEMS (GWAS Effect Mechanisms via Sequence-to-Function), a framework that trains supervised classifiers on multimodal Borzoi VEPs (ATAC-seq and RNA-seq) to distinguish fine-mapped GWAS variants from nearby (*±*500*kb*) controls, reasoning that cell types whose predictions are most discriminative should reflect disease-relevant regulatory programs. SHAP-based attributions from the trained classifiers assign individual GWAS variants to the cell types driving their predicted disease-association status, resolving the cellular context through which individual risk variants act. Applied to four diseases with well-characterized cell-type etiologies (asthma, LDL cholesterol, multiple sclerosis, and type 2 diabetes), GEMS recovered known disease-relevant cell types at both the trait and individual-variant level and linked variant-level associations to specific transcription factor disruptions, demonstrating that integrative analysis of S2F predictions can resolve cellular disease mechanisms from aggregate enrichment down to single-variant regulatory logic.

## RESULTS

### Framework overview

GEMS dissects the disease-relevant cellular mechanisms underlying GWAS signals by leveraging epigenomic (ATAC-seq) and transcriptomic (RNA-seq) variant effect predictions (VEPs) from sequence-to-function models. For a given trait, Random Forest classifiers are trained to distinguish fine-mapped causal variants (posterior inclusion probability, PIP *>* 0.1) from control variants (PIP *<* 0.01) sampled from the surrounding *±*500*kb* window, using cell-type-specific Borzoi VEPs as predictive features. In order to understand locus-level effects in terms of specific variants, SHAP analysis of the trained classifier then associates individual GWAS variants with disease-specific cellular contexts (Methods)(Figure 1).

**Figure 1.**
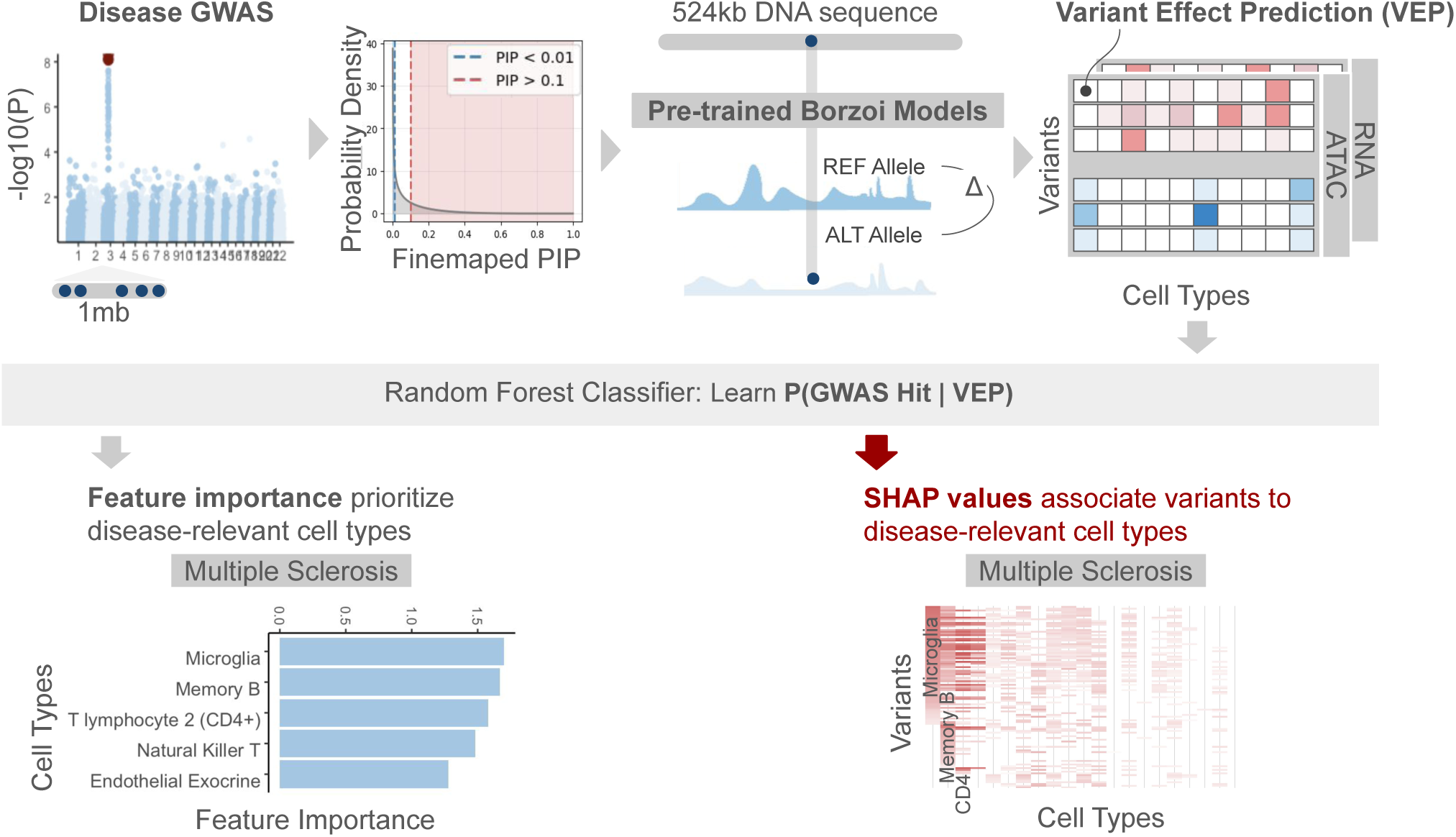
GEMS framework for mapping GWAS variants to causal cell types via sequence-to-function modeling. Fine-mapped GWAS variants (PIP *>* 0.1) and locally matched control variants (PIP *<* 0.01, sampled from *±*500*kb* windows) are input to pre-trained Borzoi sequence-to-function models, which predict reference and alternative allele effects across hundreds of cell-type-specific molecular tracks (RNA-seq and ATAC-seq). The resulting variant effect prediction (VEP) matrix (variants *×* cell types) is used to train Random Forest classifiers that distinguish causal from control variants. GEMS prioritized disease-relevant cell types (left; multiple sclerosis shown as an example), and reported variant-level cell-type associations, derived from per-variant SHAP attributions (right).

We applied GEMS to European-ancestry GWAS of four phenotypes with well-established cell-type etiologies: asthma (ASM; UK Biobank, *N* = 361,141) and LDL cholesterol (LDL; UK Biobank, *N* = 344,613)^22,23^, multiple sclerosis (MS; International Multiple Sclerosis Genetics Consortium, *N* = 68,374)^24^, and type 2 diabetes (T2D; Xue et al., *N* = 659,316)^25^. Genome-wide SuSiE fine-mapping of the GWAS summary statistics for each trait (Methods, Supplementary Table S1) identified 547, 985, 273, and 497 candidate causal variants (PIP *>* 0.1) for ASM, LDL, MS, and T2D, respectively. Fine-mapping was performed with standard SuSiE-RSS and uniform priors (Methods) to avoid circularity between functionally informed variants prioritization and regulatory signals captured by Borzoi predictions. For each causal variant, we randomly sampled, without replacement, five control variants from all variants within *±*500 kb with PIP *<* 0.01. Borzoi VEPs were computed for all variants across primary adult tissues, excluding fetal, embryonic, cell-line, and perturbation-derived specimens (Methods, Supplementary Table S2). As genetic effects on accessibility and gene expression are known to act across different scales of distance^10,26,27^, window sizes for VEPs in ATAC– and RNA-seq were chosen separately and to maximize averaged 5-fold cross validation predictive accuracy (Methods, Supplementary Figure S1). Following quality control and feature filtering (Methods), the final feature set comprised 111 ATAC-seq and 160 RNA-seq tracks (Supplementary Table S2).

For each trait, we trained Random Forest classifiers to distinguish causal from control variants using VEPs as features, fitting three independent models per trait: ATAC-seq only, RNA-seq only, and a multimodal model using predictions in both modalities. To accommodate the stochasticity of random forest models, we trained 100 replication models (same data matrix, different random state) and 100 permutation models on randomly shuffled datasets (different label-shuffled data matrix, different random state) for each trait. A hierarchical model selection criterion was used during hyperparameter tuning to jointly optimize predictive accuracy and biological specificity: hyperparameter configurations that yield the top-level averaged held-out accuracy were retained, and the final model was selected as that maximizing the ‘informativity score’, which is defined as the separation of true cell type feature importances from replication models versus a null distribution from the permutation models (Methods, Supplementary Figure S2). Individual GWAS variants were linked to causal cell types where SHAP values computed across 100 replication models were significantly higher than those from the 100 permutation models with FDR-corrected *p <* 0.05.

In the sections that follow, we evaluate GEMS at three levels. First, we assess whether Borzoi variant effect predictions discriminate fine-mapped GWAS variants from matched controls and whether the most predictive cell types recover disease-relevant cell types. Second, we resolve variant-level cell-type associations and validate them against independently characterized cis-regulatory elements (cRE). Finally, we characterize the transcription factor programs disrupted by disease-associated variants, linking cell-type associations to regulatory mechanisms at individual loci and across variant subgroups.

### GEMS recapitulate disease-relevant cell-type specificity across traits and modalities

We first asked whether Borzoi sequence-to-function predictions carry sufficient information about variant regulatory impact to distinguish fine-mapped GWAS variants from nearby non-causal controls. Across the four traits and three modalities, RF classifiers trained on VEPs discriminated causal variants from background controls (median AUROC = 0.63, range: 0.58–0.68, Supplementary Figure S3). AUPRC fold-enrichment over the prevalence-matched baseline ranged from 1.2 (T2D) to 1.6 (MS, Figure 2A), with the multimodal model achieving the highest enrichment across traits. Although single-cell ATAC-seq has been the predominant modality for annotating GWAS variants with cellular context, RNA-seq features consistently outperformed chromatin accessibility alone across all traits (mean ΔAUPRC = +0.03), likely reflecting the broader diversity of cell and tissue types represented in Borzoi’s RNA-seq tracks (160 tracks versus 111 ATAC-seq tracks) and the greater sequencing depth of bulk RNA-seq relative to single-nucleus ATAC-seq pseudobulk profiles. The multimodal model added a further gain for ASM (AUPRC fold-enrichment from 1.39 to 1.43) and MS (from 1.49 to 1.57), suggesting the two modalities carry partially non-redundant signal.

**Figure 2.**
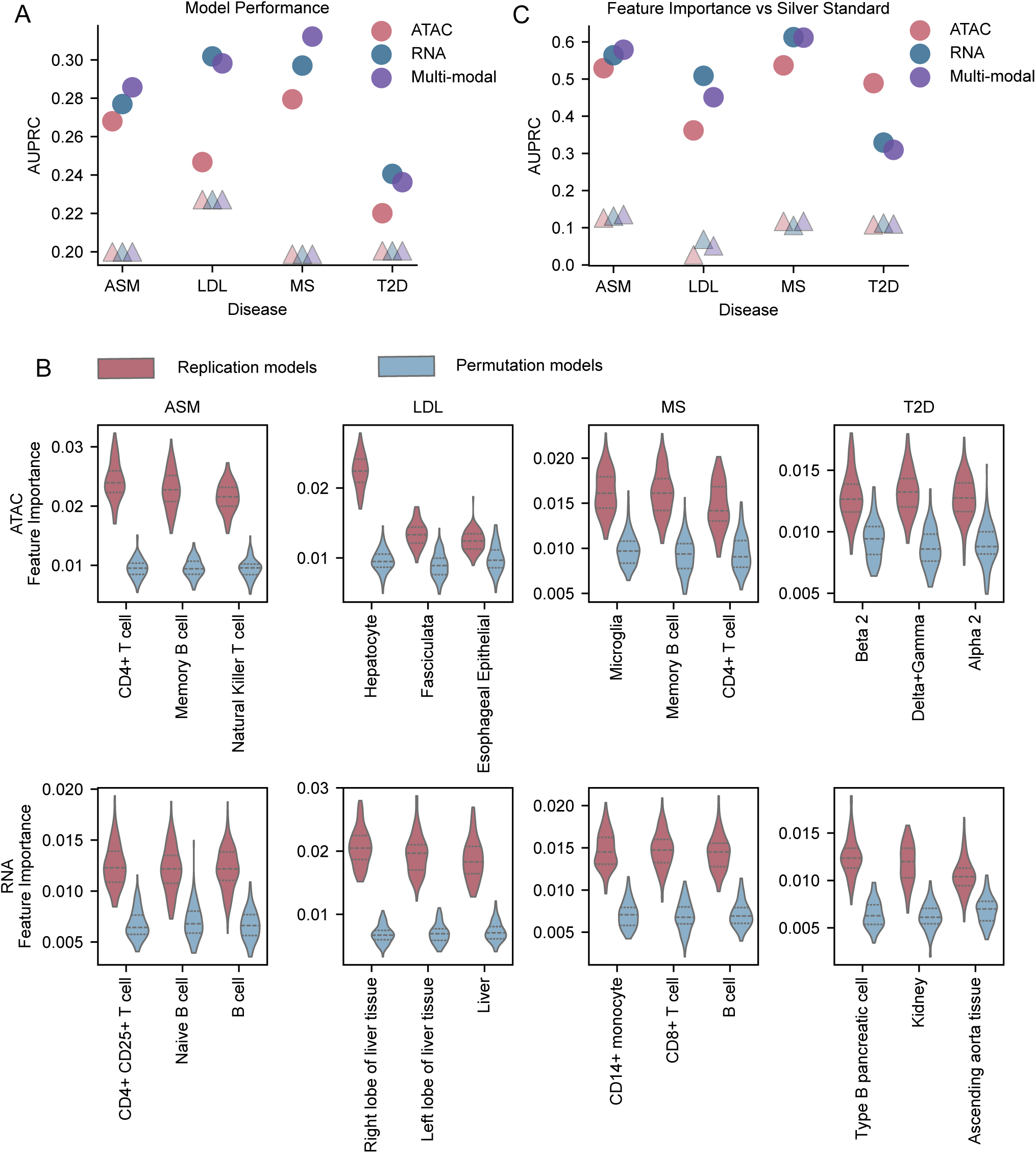
GEMS recapitulate disease-relevant cell-type specificity across traits and modalities. **(A)** AUPRC of Random Forest classifiers trained to distinguish fine-mapped GWAS variants (PIP *>* 0.1) from distance-matched controls across four diseases (ASM, LDL, MS, T2D) and three modalities (ATAC, RNA, multimodal). Circles indicate model AUPRC; triangles indicate the trait-specific random baseline (proportion of positives). **(B)** Top three cell-type features ranked by mean Random Forest importance across 100 replicates (red) compared with 100 label-permuted null models (blue) for each disease, shown separately for ATAC-seq (top row) and RNA-seq (bottom row). **(C)** Feature importance AUPRC evaluated against the silver-standard set of disease-relevant cell types for each trait and modality. Triangles indicate baseline prevalence of silver-standard cell types among all features.

Classifier performance alone does not guarantee that the most predictive features correspond to disease biology rather than correlated technical signal. We next examined whether the specific cell types driving classifier predictions reflect genuine biological signal. For each trait and modality, the feature importance values of the most predictive cell types from the replicate models showed separation from the null distributions derived from the permutation models (Figure 2B; Methods). Across all four diseases, the top-ranked cell types aligned with established causal biology: CD4^+^ T lymphocytes (ATAC and RNA) for ASM, hepatocytes (ATAC) and liver (RNA) for LDL, memory B cells (ATAC and RNA) and microglia (ATAC) for MS, and pancreatic islet subtypes (ATAC: alpha 2, beta 2, delta+gamma) and pancreas cells (RNA) for T2D (Figure 2B; Supplementary Table S3). While Borzoi RNA-seq feature set does not include a microglia-specific track, the RNA model identified CD14^+^ monocytes as the most predictive feature for MS. Despite their distinct ontogenetic origins, circulating monocytes share extensive transcriptomic homology and functional similarities with microglia, making them a robust transcriptomic proxy for CNS-resident myeloid lineages^28–30^.

To quantify whether disease-relevant cell types are systematically more predictive, we required a reference list of causal cell types for each trait. Because no comprehensive, experimentally validated gold-standard catalogue of all causal cell types exists for any complex disease, we instead curated a “silver standard”: a set of disease-relevant cell types derived from published literature via AI-assisted search followed by manual review (Methods). Across our focal traits, a median of 11.3% (range: 2.7–12.6%) of ATAC features and 10.9% (range: 6.9–13.1%) of RNA features are included in the silver standard (Supplementary Table S4). Ranking cell types by feature importance and evaluating against the silver standard yielded high discriminative accuracy, with AUROCs ranging from 0.66 (LDL, ATAC) to 0.91 (MS, RNA, Supplementary Figure S4) and AUPRCs showing a 2.8– to 13.4-fold enrichment over the trait-specific baseline prevalences (Figure 2C; Table S5). LDL had the lowest AUROC (0.66), reflecting the class imbalance in silver standard for this trait (3 of 111 ATAC cell types are labeled as “silver standard”, while the rest are not, 2.7%, an extreme imbalance). Nevertheless, the GEMS ATAC model ranked hepatocytes as the top feature, consistent with the established role of hepatocellular LDLR-mediated cholesterol clearance.

Taken together, these results established that Borzoi VEPs encode cell-type-specific regulatory information sufficient to recover disease-relevant biology at the trait level.

### GEMS recovers disease-relevant cell types concordant with sLDSC

To evaluate GEMS-prioritized cell-types against an orthogonal, sequence-free method, we compared GEMS cell-type significance (see Methods) with coefficient *z*-scores from stratified LD-score regression (sLDSC), which quantifies regulatory enrichment directly from GWAS summary statistics. To ensure a fair comparison, we ran joint sLDSC using CATlas snATAC-seq peak annotations for the same 111 cell types used by GEMS (ATAC).

The two methods converged on the same disease-relevant cell types in most cases. Both GEMS and sLDSC ranked CD4^+^ T lymphocytes and hepatocytes at the top for ASM and LDL, respectively (Figure 3A). For T2D, both methods converged on pancreatic islet subtypes as the dominant signal, though the top-ranked subtype differed: sLDSC prioritized beta 1, while GEMS ranked delta+gamma, alpha 2, and beta 2 as the top three. For MS, both methods identified memory B cells and T lymphocytes among the top-ranked cell types (sLDSC: CD8^+^ T, memory B; GEMS: microglia, memory B, NKT, CD4^+^ T). sLDSC did not detect the microglial signal captured by GEMS (ATAC), consistent with the lower yield and poorer peak coverage of microglia in single-cell atlases^31^ that would attenuate annotation-based enrichment signals.

**Figure 3.**
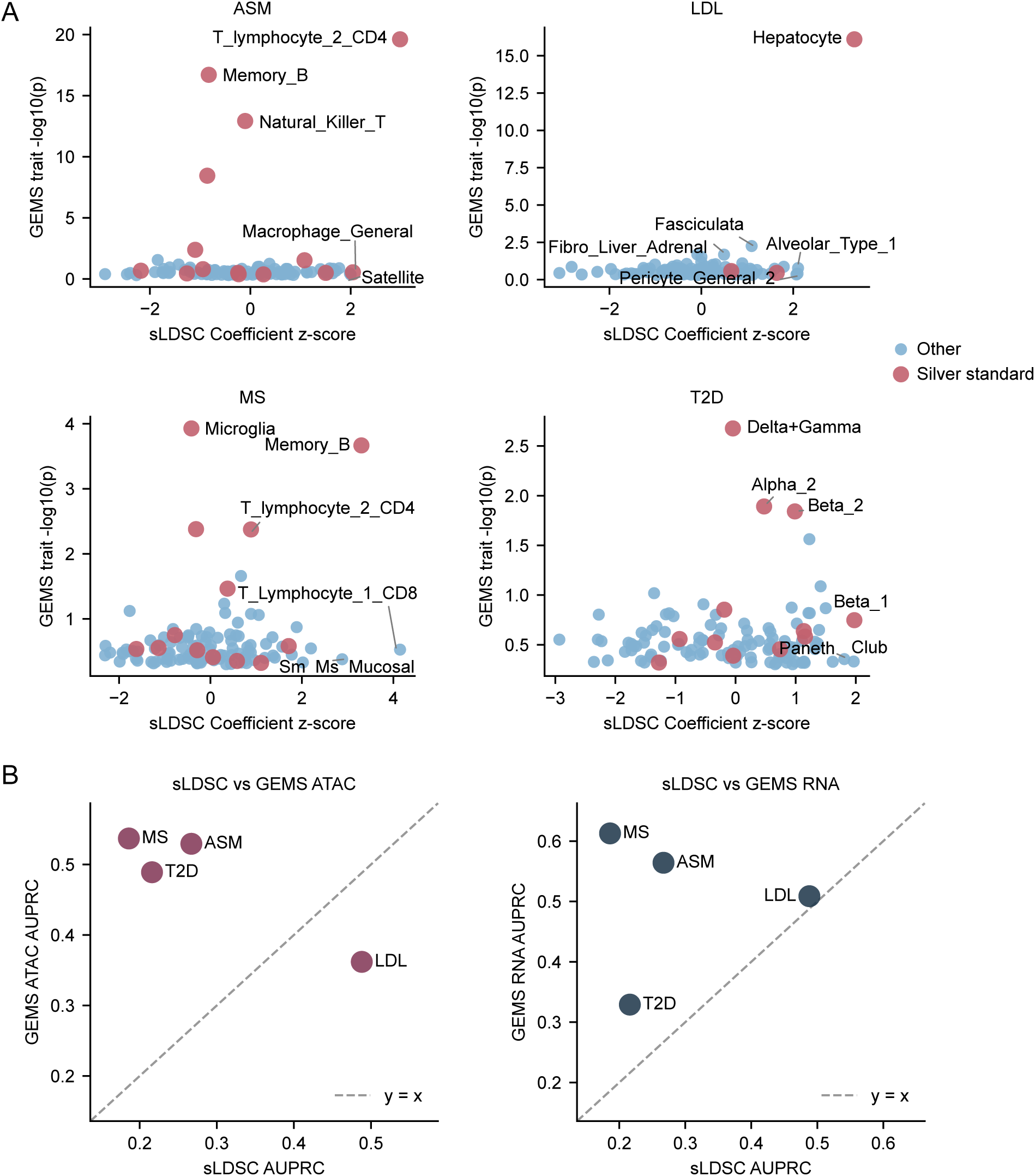
GEMS recovers disease-relevant cell types concordant with sLDSC. **(A)** GEMS cell-type significance (*−* log_10_ *P*) versus sLDSC coefficient *z*-scores for 111 CATlas snATAC-seq cell types across four diseases. Each point represents a cell type; pink points denote silver-standard cell types, blue points denote remaining cell types. The top three cell types by each method (GEMS and sLDSC) are labeled. **(B)** Silver-standard AUPRC for GEMS versus sLDSC across four traits, shown separately for ATAC (left) and RNA (right). Dashed line indicates *y* = *x*; points above the line indicate higher GEMS AUPRC than sLDSC.

To compare the overall ability of GEMS and sLDSC to prioritize disease-relevant cell types, we calculated the AURPC for identifying silver-standard cell types using rankings based on sLDSC coefficient *z*-scores or GEMS cell-type significance. GEMS achieved higher AUPRC than sLDSC for three of four traits (ASM, T2D, and MS) using ATAC-seq features and for all four traits using RNA-seq features, with median AUPRC gains of 0.27 and 0.21, respectively (Figure 3B). LDL was the exception in the ATAC comparison, again reflecting the imbalanced silver-standard class distribution for this trait.

These analyses demonstrate that GEMS independently recovers the same disease-relevant cell types detected by sLDSC heritability enrichment, with higher overall silver-standard enrichment in most trait-modality combinations. Together with the classifier and silver-standard evaluations above, these trait-level analyses provide the foundation for variant-level cell-type associations in the following sections.

### GEMS resolves variant-level cell-type associations across diseases and modalities

To assign variant-level cell-type associations, we computed SHAP values for each GWAS variant across all cell-type tracks from the 100 replication models for the corresponding GWAS and tested them against a null distribution derived from permutation models (see Methods). Variant–cell-type pairs with *FDR <* 0.05 (one-tailed Z-test, Benjamini–Hochberg correction per variant) were retained as a significant association.

Across the four traits, GEMS associated a median of 65.9% (range: 54.2–75.5%) fine-mapped GWAS variants with at least one cell type using RNA-seq features and a median of 58.7% (range: 49.8–67.8%) using ATAC-seq (Table S6). These variant–cell-type associations were significantly enriched for the corresponding trait– and modality-specific silver-standard cell types (median fold enrichment = 4.8, binomial test p-value *<* 2.2 *×* 10*^−^*^16^, see METHOD). By comparison, only 26% of fine-mapped GWAS variants overlapped a CATlas snATAC-seq peak for any cell type, indicating that GEMS extends cell-type annotation to the majority of variants that lack detectable chromatin accessibility peaks in existing single-cell atlases.

For most of the trait-modality combinations, a majority (median of 55.0%) of significant associations mapped to a single dominant cell type that matched the most well-established disease-relevant biology (Table S7). For T2D, 51.5% (256/497) of fine-mapped variants (*PIP >* 0.1) were associated with bulk pancreatic tissue by RNA-seq (Figure 4A). While the RNA-seq features lack cell-type resolution, snATAC-seq resolved 26.0% (129/497) of the T2D GWAS variants to pancreatic islet cell-types (alpha_2_, beta_2_, and delta+gamma; Figure S5), consistent with the predominant role of pancreatic islet dysfunction in T2D pathogenesis^32,33^. The lower proportion associated through snATAC-seq may reflect the underrepresentation of pancreatic islet cells in CATlas. For ASM, CD4^+^ T lymphocytes^34,35^ were the most commonly associated cell type in both modalities (34.7% in ATAC, 34.0% in RNA, Figure S5). LDL variants converged on hepatocytes (45.7% in ATAC) and liver tissue (40.0% in RNA), reflecting the central role of hepatocellular *LDLR*-mediated cholesterol clearance^36,37^. For MS, 31.9% of variants were associated with memory b cells^38,39^ and microglia^40,41^ by ATAC-seq and 37.4% with CD14+ monocytes in RNA-seq.

**Figure 4.**
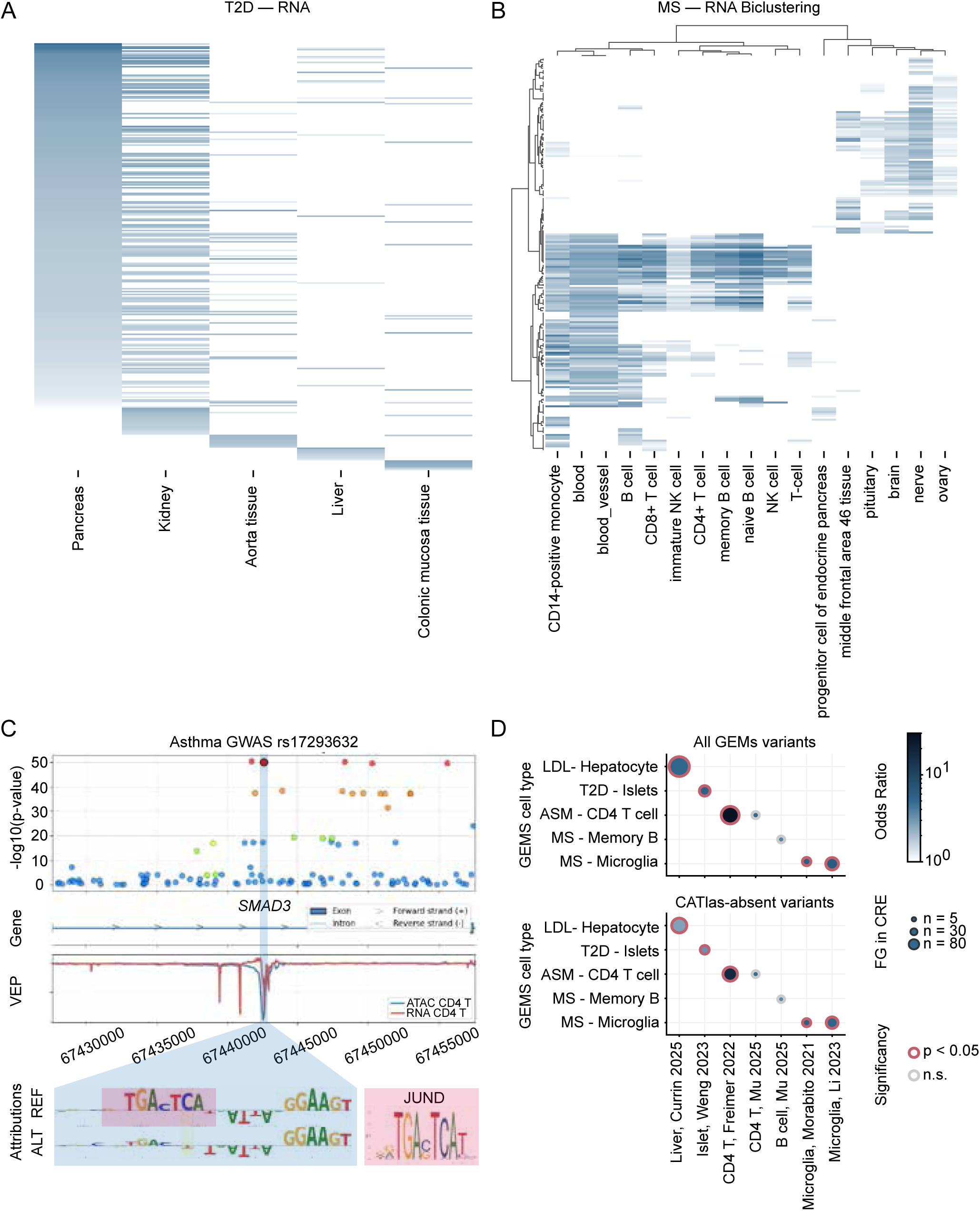
GEMS resolves variant-level cell-type associations and validates them against independent cis-regulatory elements. **(A)** Variant–cell-type association heatmap for T2D RNA-seq. **(B)** Hierarchical biclustering of MS RNA-seq variant–tissue association matrix revealing immune and CNS variant modules. **(C)** Three fine-mapped variants in a single asthma credible set at the *SMAD3* locus. Top: LocusZoom plots; middle: Borzoi VEP profiles in CD4^+^ T cells; bottom: TF motif disruption analysis showing JunD (AP-1) motif disrupted by rs17293632 C*>*T. **(D)** Dot plot of Fisher’s exact test enrichment for GEMS-associated variants in cell-type-matched cREs from independent studies. Top: all GEMS-associated variants; bottom: CATlas-absent variants only. Dot color indicates odds ratio (log scale); dot size indicates foreground variants overlapping cREs; red border indicates *P <* 0.05.

Beyond the immune associations, 25.2% of MS variants were associated with bulk brain tissue by the RNA-seq model, recapitulating the well-established role of neuronal dysfunction in MS pathogenesis^42,43^. Hierarchical biclustering of the MS RNA-seq variant–tissue association matrix revealed two principal variant modules: one (114 variants) driven by peripheral immune cells such as monocytes, B cells, and lymphocytes, and a second (92 variants) enriched for brain and nerve tissues (Figure 4B), partitioning MS risk variants into two discrete modules with minimal cross-assignment.

To determine whether these two variant modules operate through distinct regulatory programs, we applied input *×* gradient attibution and TF-MoDISco motif discovery to identify TF binding motifs disrupted in each variant-cell-type associations (see METHOD). We tested whether specific TF families were preferentially disrupted in each diagonal block of the biclustering (immune variants *×* immune tissue, or CNS variants *×* CNS tissues) relative to the remainder of the heatmap using Fisher’s exact tests with BH correction (Table 1). The Ets-related family was the most significant enriched in the immune*×*immune block (OR = 3.5, FDR = 1.1 *×* 10*^−^*^5^), driven primarily by *SPIB*, *ELF4*, and *ELF3*. The ETS TF family is a crucial master regulator of immune cell fate^44^ and a major driver of genetic susceptibility^45^ to MS. Additional families enriched in the immune*×*immune block included STAT^46^ (OR = 5.6, FDR = 0.04) and C/EBP-related^47^ (OR = 4.7, FDR = 0.048), both of which encode regulators of myeloid and lymphoid differentiation^48^. In contrast, the CNS*×*CNS block was dominated by the Tal-related family (OR = 24.6, FDR = 1.5 *×* 10*^−^*^7^), driven by *NGN2* and *MSGN1*, implicating disruption of neuronal differentiation programs as a distinct mechanism from the immune-mediated pathology captured by the first module^49,50^.

**Table 1.** Enrichment of GEMS-associated variants not overlapping CATlas ATAC-seq peaks. FG: foreground variants after removing those overlapping CATlas ATAC-seq peaks for the matched cell type; FG total: total CATlas-absent foreground size. Background is unchanged from Table S9.

| Disease | Cell type | CRE source | FG in cRE | FG total | BG in cRE | BG total | OR | P |
| --- | --- | --- | --- | --- | --- | --- | --- | --- |
| ASM | CD4<sup>+</sup> T | Freimer et al. | 49 | 155 | 9 | 361 | 18.08 | $5.7 \times 10^{-20}$ |
| ASM | CD4<sup>+</sup> T | Mu et al. | 4 | 155 | 2 | 361 | 4.76 | $7.0 \times 10^{-2}$ |
| LDL | Hepatocyte | Currin et al. | 54 | 393 | 30 | 545 | 2.73 | $1.2 \times 10^{-5}$ |
| T2D | Islet (all) | Weng et al. | 12 | 117 | 13 | 376 | 3.19 | $5.7 \times 10^{-3}$ |
| MS | Memory B | Mu et al. | 3 | 59 | 3 | 204 | 3.59 | $1.3 \times 10^{-1}$ |
| MS | Microglia | Morabito et al. | 9 | 54 | 7 | 219 | 6.06 | $9.3 \times 10^{-4}$ |
| MS | Microglia | Li et al. | 29 | 54 | 39 | 219 | 5.35 | $2.6 \times 10^{-7}$ |

Taken together, these results demonstrate that GEMS associates the majority of fine-mapped GWAS variants with disease-relevant cell types and, when combined with TF motif analysis, can further decompose polygenic risk into mechanistically distinct regulatory programs, which we next apply to resolve individual causal variants within unresolved credible sets.

### Variant-level resolution at the *SMAD3* asthma locus

Variant-level cell-type associations provide an opportunity to investigate the regulatory mechanism of individual GWAS variants. We examined a three-variant credible set located within the first intron of gene *SMAD3* where fine-mapping could not confidently nominate a single causal variant: rs72743461 (PIP = 0.37), rs56375023 (PIP = 0.26), and rs17293632 (PIP = 0.21) all reached genome-wide significance with nearly identical association statistics (Figure 4C, top). GEMS associated rs17293632 with CD4^+^ T lymphocytes in both the ATAC-seq and RNA-seq models, with Borzoi VEPs showing a strong reduction in predicted chromatin accessibility (log_2_ FC = *−*0.24) and a concordant decrease in predicted *SMAD3* expression (log_2_ FC = *−*0.08, Figure 4C, middle). In contrast, rs72743461 showed a significant association with CD4^+^ T cells in ATAC-seq but negligible predicted effect (log_2_ FC = 0.01) and no RNA-seq support, while rs56375023 showed RNA-seq association but weak VEP (log_2_ FC = 0.01) and no ATAC-seq signal.

To identify the molecular mechanism underlying the rs17293632 signal, we performed TF motif analysis on the CD4^+^ T cell ATAC-seq and RNA-seq output tracks (Methods). In both modalities, the C*>*T substitution was predicted to disrupt a JunD binding motif (AP-1 family, Figure 4C, bottom), consistent with the predicted loss of local chromatin accessibility and *SMAD3* expression (Figure 4C, middle). Elevated level of *SMAD3* has been reported to drive severe airway wall thickening and excessive smooth muscle contraction^51,52^. The predicted direction of effect for the T allele is consistent with its protective association with asthma (GWAS *β* = *−*0.012). The disruption of AP-1 binding by the rs17293632 T allele has been experimentally validated by Turner et al.^53^. In contrast, rs56375023 showed no detectable motif disruption, and rs72743461 was predicted to disrupt a ZIC5 motif with no established role in T cell regulation and with weak attribution signal (12-fold lower than the rs17293632 JunD attribution).

Together, the convergence of GEMS cell-type association, Borzoi VEP magnitude, and TF motif disruption across both modalities identifies rs17293632 as the most likely functional variant at this locus. This variant has a well-established genetic association with asthma^54,55^, and its disruption of AP-1 binding has been experimentally validated in cardiovascular disease models^53^.

### GEMS variant–cell-type associations are enriched in experimentally validated cis-regulatory elements

To verify whether GEMS variant–cell-type associations reflect genuine regulatory biology, we assessed their overlap with independently characterized cis-regulatory elements (cREs) from experimental datasets not used to train Borzoi. For each disease, we curated cRE sets from six published studies^56–61^ targeting the cell type most frequently associated by GEMS using ATAC-seq (Methods, Supplementary Table S8). We then tested whether GWAS variants assigned to that cell type (foreground) were enriched for cRE overlap relative to the remaining fine-mapped GWAS variants (background) using one-sided Fisher’s exact tests. For MS, we additionally tested microglia, the second most frequently associated cell type, given its well-established causal role in MS pathogenesis.

GEMS-associated variants were significantly enriched in cell-type-matched cREs for all four diseases, with odds ratios ranging from 3.7 to 28.7 across all seven tests (median 6.1 across LDL-Hepatocyte-Currin^56^, T2D-Pancreatic islet-Weng^57^, ASM-CD4 T cell-Freimer^58^, ASM-CD4 T cell-Mu^59^, MS-Memory B cell-Mu^59^, MS-Microglia-Morabito^60^, MS-Microglia-Li^61^; Figure 4D, top; Table S9). The strongest enrichment was observed for ASM CD4^+^ T cell variants in Freimer et al.^58^ ATAC-seq peaks, which also comprised one of the largest cRE sets tested (236,975 peaks, OR = 28.7, *P* = 5.3 *×* 10*^−^*^33^). The two tests that did not reach nominal significance used comparatively small cRE datasets^59^ (36,235 peaks for CD4^+^ T cells and 36,192 for memory B), but showed directionally consistent odds ratios (OR = 3.8 and 4.9 respectively; Table S9).

A substantial fraction (78.7%-92.1%) of GEMS-associated variants do not overlap a CATlas ATAC-seq peak for the matched cell type, raising the question of whether these associations reflect regulatory sequence patterns beyond those captured by the chromatin accessibility data used to train Borzoi. To test this, we repeated the cRE enrichment analysis after restricting the foreground to variants without a CATlas peak overlap for the matched cell type (Table S9). Significant enrichment persisted for all disease–cell-type pairs that reached significance in the full analysis (Figure 4D, bottom; Table 1), with attenuated but still substantial odds ratios (range 2.7-10.1, median 4.8), indicating that the regulatory sequence patterns learned by Borzoi generalize to variants outside the chromatin accessibility peaks used for training.

Having validated ATAC-seq associations against experimentally characterized cREs, we used a complementary strategy to assess the biological relevance of RNA-seq associations. Because RNA-seq associations reflect predicted transcriptional effects rather than chromatin accessibility, an analogous cRE overlap test is not applicable. Instead, we asked whether variants that the RNA model assigns to the most disease-relevant cell type tend to lie near genes in disease-relevant biological pathways. For each disease, we identified protein-coding genes within the Borzoi RNA prediction window (31,622 bp) of variants associated with the dominant cell type (foreground) and compared their pathway composition to that of genes near variants assigned to any other cell type (background), using one-sided Fisher’s exact tests (Methods).

LDL showed the strongest pathway specificity, with foreground genes near liver-associated variants significantly enriched in the cholesterol homeostasis pathway^37,62^ (OR = 6.9, *P* = 0.019; Table S10). For the remaining diseases, disease-relevant pathways showed directionally consistent odds ratios but did not reach statistical significance, likely reflecting small background gene sets (39–124 genes): T cell differentiation for ASM (OR = 3.9), regulation of NIK/NF-*κ*B signaling for MS (OR = *∞*, 4 versus 0 background genes), and glucose homeostasis for T2D (OR = 2.5) (Table S10).

### Multimodal analysis identifies variants with concordant chromatin and transcriptional associations

Having evaluated ATAC-seq and RNA-seq variant associations independently, we next asked whether individual variants show concordant associations across both modalities in the multi-modal GEMS model, which would provide convergent evidence that a variant simultaneously disrupts chromatin accessibility and gene expression in the same disease-relevant cell type. Using SHAP values from the multimodal model, we identified variants with at least one significant ATAC-seq and at least one significant RNA-seq association (FDR *<* 0.05). For each disease, we ranked (RNA-tissue, ATAC-cell-type) pairs by the number of shared variants.

Cross-modal concordance was substantial in three of four diseases. A total of 167 of 547 ASM variants (30.5%), 191 of 985 LDL variants (19.4%), and 80 of 273 MS variants (29.3%) carried significant associations in both modalities (Table 2), where the top-ranked cross-modal pairs showed significantly non-random overlap (hypergeometric test) and recapitulated disease-specific biology: liver RNA-seq with hepatocyte ATAC-seq for LDL (149 of 191 multimodal variants, 78.0%; 1.7-fold enrichment, *P* = 4.5 *×* 10*^−^*^22^), CD4^+^ T lymphocyte in both modalities for ASM (100/167, 59.9%; 2.2-fold, *P* = 1.7 *×* 10*^−^*^28^), and B cells in RNA-seq with memory B cell ATAC-seq for MS (50/80, 62.5%; 2.8-fold, *P* = 1.7 *×* 10*^−^*^23^; Table 2).

**Table 2.**
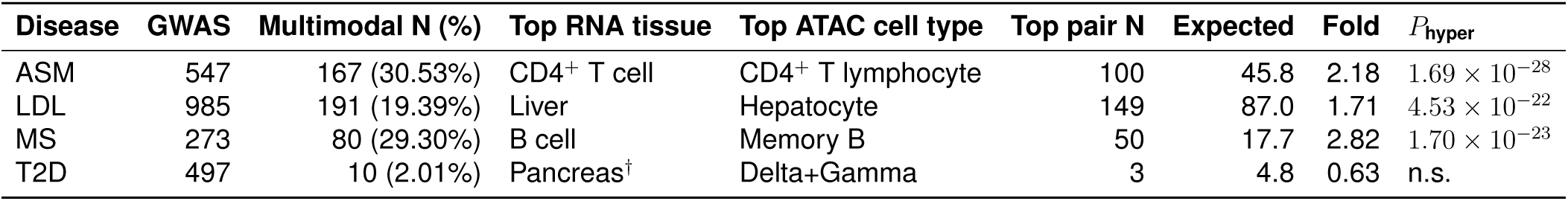
Cross-modal variant–cell-type concordance in the multimodal model. GWAS: finemapped variants with PIP *>* 0.1. Multimodal N: variants with at least one significant ATAC-seq and one significant RNA-seq association (FDR *<* 0.05) in the multimodal RF model, after excluding mononuclear cell from RNA features. Top pair: the (RNA-tissue, ATAC-cell-type) pair shared by the most variants. Expected: expected overlap under independent assortment of ATAC and RNA associations. Fold: observed/expected ratio. *P*_hyper_: one-sided hypergeometric test. *^†^*Tied with colon and colonic mucosa (3 variants each).

T2D was a notable outlier: despite substantial single-modality association rates (26.0% ATAC, 51.5% RNA; Table S7), only 10 of 497 variants (2.0%) showed cross-modal concordance in the multimodal model. This was primarily driven by far fewer significant ATAC-seq cell-type associations in the multimodal model: only 21 of 497 T2D variants (4.2%) were significantly associated with any ATAC-seq cell type, compared with 31.0–41.0% in the other three diseases, whereas the RNA-seq association rate (63.4%) was comparable to other phenotypes (52.2–69.5%). Consistent with this, the T2D ATAC-only model showed the weakest improvement over baseline for discriminating causal GWAS variants (AUPRC fold-change 1.1; Figure 2A). The lower proportion of variants associated using snATAC-seq may reflect limited representation of pancreatic islet cell types in CATlas. The islet ATAC-seq pseudobulk tracks were constructed from approximately 15,000 nuclei^11,63^, with rarer endocrine subtypes, such as delta+gamma cells, represented by only approximately 710 nuclei. This limited representation may reduce peak coverage and sensitivity for detecting variant associations in these cell types

## DISCUSSION

The analyses presented here demonstrate that supervised meta-modeling of sequence-to-function variant effect predictions can resolve the cellular context of GWAS variants at both the trait and individual-variant level. Across four diseases with well-characterized cell-type etiologies, GEMS recovered known disease-relevant cell types from aggregate feature importances, associated the majority of fine-mapped GWAS variants with at least one cell type, and linked variant-level associations to specific transcription factor programs. These associations extended to variants lacking detectable chromatin accessibility peaks in the single-cell atlas data used to train Borzoi, and were validated against cis-regulatory elements identified by independent experimental studies.

A key challenge in applying supervised classifiers to variant effect predictions is that Borzoi outputs across cell types are highly correlated and the discriminative signal between causal and control variants is modest. Under these conditions, random forest classifiers can achieve similar predictive accuracy while relying on entirely different sets of cell types across hyperparameter configurations or random seeds^64^, making model selection on accuracy alone an unreliable guide to biological interpretation. To address this, we defined an informativity score that measures the separation of a model’s cell-type feature importances from a permutation-based null distribution (Methods). Among models with comparable held-out accuracy, those with higher informativity scores consistently prioritized known disease-relevant cell types, and a hierarchical selection strategy that jointly optimized accuracy and informativity recovered biologically interpretable results across all four diseases.

These considerations motivated a comparison with stratified LD-score regression (sLDSC), which quantifies heritability enrichment directly from GWAS summary statistics rather than through supervised classification of variant effect predictions. The cell types most predictive in GEMS were broadly concordant with those showing the highest heritability enrichment in sLDSC when both methods operated on the same CATlas snATAC-seq annotations. GEMS provided additional resolution for phenotypes where sLDSC is limited by annotation sparsity or feature correlation: for multiple sclerosis, GEMS identified microglia and memory B cells as the most relevant cell types, whereas sLDSC ranked CD8^+^ T lymphocytes highest, consistent with the difficulty of resolving cell types with sparser chromatin accessibility peaks and high inter-cell-type correlation among immune populations (Figure S9). These results suggest that the supervised, prediction-based approach of GEMS can complement enrichment-based methods by leveraging the regulatory grammar learned by sequence-to-function models rather than relying solely on peak-level overlap.

A notable feature of these data is the consistent superiority of RNA-seq over ATAC-seq variant effect predictions, both in discriminating fine-mapped GWAS variants from controls (Figure 2A) and in resolving variant-level cell-type associations (Table S7). Although single-cell ATAC-seq has been the predominant modality for annotating GWAS variants with cellular context^11,12^, RNA-seq features outperformed chromatin accessibility across all four diseases. This asymmetry may reflect differences in the training data underlying each modality: Borzoi’s ATAC-seq tracks were trained on single-nucleus ATAC-seq pseudobulk profiles from the CATlas atlas^63^, which are inherently sparser than the bulk RNA-seq data from ENCODE and GTEx that underpin the transcriptomic tracks. The multimodal model provided a further gain for asthma and multiple sclerosis, indicating that the two modalities carry partially non-redundant regulatory signal, but the marginal contribution of ATAC-seq was modest relative to RNA-seq alone.

These observations extend from the trait level to individual variants. GEMS associated more than half of fine-mapped GWAS variants with at least one cell type, providing cellular context for variants that would otherwise lack annotation through overlap-based methods using the same underlying experimental data. Variants that did not overlap CATlas ATAC-seq peaks for their associated cell type were still significantly enriched in cis-regulatory elements from independent experiments targeting that cell type (Table 1, Table S9), indicating that Borzoi captures regulatory sequence features beyond those reflected in observed chromatin accessibility peaks. Collectively, these findings are consistent with a model in which sequence-to-function predictions encode latent regulatory information that can be surfaced through supervised classification even at loci where experimental chromatin profiling data are sparse or absent.

We acknowledge several limitations. First, sequence-to-function models have difficulty distinguishing cell types with similar regulatory programs, and GEMS inherits this constraint: variants are typically associated with multiple functionally related cell types rather than a single causal cell type. Second, the repertoire of cell types available for analysis is constrained by the experimental data used to train the underlying model. Tissues that are difficult to obtain, such as pancreatic islets, are underrepresented in training corpora, and GEMS has correspondingly limited power in those cellular contexts, as evidenced by the weak ATAC-seq signal for T2D. Third, much of the RNA-seq data underlying Borzoi derives from bulk tissue profiling rather than single-cell assays, limiting resolution to the cell-type or cell states. Fourth, GEMS requires a sufficient number of fine-mapped GWAS variants to train the classifier, restricting application to well-powered GWAS. Rare diseases and historically underpowered studies may not provide enough training signal. Finally, both GEMS and sLDSC showed reduced performance when applied to alternative GWAS studies of the same disease (Figure 5A), indicating that study-level characteristics—cohort composition, phenotype definition, and statistical power distribution across loci—can influence the cell-type signals recovered by either approach.

Several aspects of this work point to productive future directions. The GEMS framework is agnostic to the source of variant effect predictions: we used Borzoi because it was the most capable sequence-to-function model available at the time of this study, but the classifier-based approach is compatible with predictions from any such model. It will be informative to assess whether next-generation models, such as the recently released AlphaGenome^19^, yield more accurate and more granular cell-type associations through this framework. The extension of GEMS to multi-ancestry GWAS, and to cellular contexts beyond those represented in current training corpora—including disease-relevant tissues obtained through emerging single-cell profiling technologies—represents a further opportunity to improve the resolution and generalizability of variant-to-cell-type mapping.

Taken together, these analyses demonstrate that integrative meta-modeling of sequence-to-function predictions can resolve the cellular mechanisms of GWAS variants, extending regulatory annotation beyond the reach of overlap-based methods and providing a framework for decomposing polygenic risk into mechanistically distinct, cell-type-specific programs.

## METHODS

### GWAS and variant set generation

We obtained GWAS summary statistics for four target phenotypes: asthma (ASM), LDL cholesterol (LDL), type 2 diabetes (T2D), and multiple sclerosis (MS). To ensure data quality, we excluded variants located in ENCODE blacklist regions (extended by a 100 bp window) and removed all insertions and deletions (indels), retaining only single-nucleotide polymorphisms (SNPs).

To define a set of likely causal variants (positives), we performed genome-wide fine-mapping with SuSiE^65^ using uniform (non-informative) priors. The genome was partitioned into overlapping 3 Mb windows (1 Mb stride), matching the pre-computed UK Biobank LD matrix blocks (*∼*380k European individuals)^66^. SuSiE was run independently in each window via sufficient statistics (susie_suff_stat), with a maximum of 10 causal signals per window, residual variance fixed at 1, and default credible set parameters (coverage = 0.95, purity *|r|*_min_ = 0.5). Before fine-mapping, DENTIST^67^ was applied to detect and remove SNPs with LD-mismatch between the GWAS summary statistics and the reference LD panel. Results were aggregated across overlapping windows by retaining each SNP’s estimate from the window in which it was most central. Variants with a Posterior Inclusion Probability (PIP) *>* 0.1 were obtained as positive variants (hits). For each identified positive variant, we sampled 5 negative variants (PIP *<* 0.01) located on the same chromosome within a *±*500 kb window (spanning 1 Mb) centered on the positive variant.

### Variant effect prediction and feature processing

We utilized the gRelu package to interface with the Borzoi deep learning model to predict the impact of variants on chromatin accessibility (ATAC-seq) and gene expression (RNA-seq). For each variant in our dataset, we generated predictions for both the reference and alternative alleles. We focused on the relevant local context for each modality, analyzing a 500 bp region centered on the variant for ATAC-seq tracks and a 31,622 bp window for RNA-seq tracks (Figure S1).

For each track, we quantified the variant effect as the absolute log2 fold change (absFC) between the alternative and reference predictions: *|* log_2_(Alt + 1) *−* log_2_(Ref + 1)*|*. These scores were standardized using z-score normalization. To ensure robust predictions, we employed an ensemble approach, averaging the outputs from four independent Borzoi model replicates.

To ensure the quality and biological relevance of the predictive features, we applied a rigorous post-processing pipeline:

- Exclusion of developmental and perturbed contexts: We filtered out tracks corresponding to fetal, embryonic, or newborn tissues, cell lines, as well as samples annotated as “treated” in ENCODE metadata for RNA-seq tracks, to focus on baseline adult physiology. We also removed mononuclear cell which is a super group of lymphocytes and monocytes.
- Strand integration (RNA-seq only): For stranded assays, we summed the predictions from the forward and reverse strands to generate a single unstranded signal.
- Replicates aggregation (RNA-seq only): Replicate tracks were averaged to produce a single consensus feature per tissue type.
- Quantile normalization (multi-modal model only): Quartile normalization was used to reduce the batch effects between ATAC-seq and RNA-seq tracks.

### Model training

We trained Random Forest classifiers to predict causal variant status using the Borzoi variant effect predictions (VEPs) as features. For each phenotype, we trained three models using different input modalities: (1) an ATAC-seq only model, (2) an RNA-seq only model, and (3) a multi-modal model integrating tracks from both ATAC-seq and RNA-seq.

We performed a hyperparameter grid search with varied the maximum tree depth (max depth: 10, 15), minimum samples per split (min samples split: 2, 5, 7, 10), minimum samples per leaf (min samples leaf: 2, 5, 7, 10), and the cost-complexity pruning parameter (ccp alpha: 0, 0.001). To address class imbalance between positive and negative variants, we applied balanced class weights during training. For each hyperparameter configuration, we performed 5-fold cross-validation.

To ensure result stability and estimate prediction variance, the full training procedure for each hyperparameter configuration was repeated 100 times using unique random seeds (replicates). To rigorously establish a baseline for feature importance, we generated a corresponding null distribution for every trained model. This was achieved by performing 100 independent training runs using the identical feature matrix but with permuted phenotype labels (null models).

Deviation from this null background serves as a quantitative measure of the true biological signal captured by the Borzoi VEPs. Feature deviation was quantified by calculating Z-scores that compared the mean importance of the real model replicates against the distribution derived from the null models. P-values were computed from these Z-scores and corrected for multiple hypothesis testing using the Benjamini-Hochberg False Discovery Rate (FDR) method.

To quantify the aggregate signal gain of each model, we defined an informativity score. This score was calculated as the sum of the negative log-transformed FDR-adjusted p-values derived from the comparison against the null distribution:

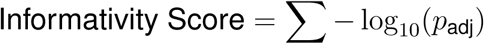

By operating in the negative log space, this transformation effectively zeros out uninformative background features. Consequently, the final informativity score is not artificially inflated by a large number of weak features, but is strictly driven by the specific features exhibiting the largest, most reliable deviation from the background.

While multiple hyperparameter configurations achieved comparable accuracy in distinguishing causal GWAS variants from non-causal baselines, they sometimes relied on bVEPs from biologically irrelevant cell types. To ensure that the selected model both maintained high performance at predicting GWAS variants and specifically leveraged bVEPs from disease-relevant cell types, we implemented a two-step hierarchical selection criterion to prioritize sensitivity and feature robustness. First, we ranked all model configurations by their accuracy on the held-out test set and retained the top 37.5% of models (Figure S10). Second, from this high-performance subset, we selected the final model that maximized the informativity score. This ensured the selection of a model with the strongest, most biologically meaningful separation from the null background.

### Disease-relevant silver standard list generation

To establish a “silver standard” set of biologically relevant cell types and tissues for each phenotype, we employed an AI-assisted literature search strategy.

We utilized OpenAI’s Deep Research tool to systematically screen the complete list of ATAC-seq and RNA-seq tracks used to train the GEMs models. For each phenotype, the model was queried with the complete track list and the following standardized prompt (using LDL cholesterol as a representative example):

> “For the phenotype Low-Density Lipoprotein (LDL), can you list cell types and tissue from this list I provided that are causal to LDL? For reference, these are the cell types and tissues from CAGE, ATAC, DNase, CHIP, RNA experiments used to train https://www.nature.com/articles/s41588-024-02053-6. For each cell type or tissue you claim to be causal, list (1) its name in my file (2) its line number in my file (3) its common name (4) some literature support (citation, paper name, paper link). Please list all possible causal items in my list, not just the unique one. And put your results in a csv file for me to download.”

The candidate lists generated by the AI model underwent a rigorous manual review process to ensure accuracy. We verified the relevance of each cited study and corrected the assigned status (“relevant” vs. “not relevant”) where necessary. This two-step process, automated AI literature search followed by manual curation, produced the final silver standard list used to benchmark model feature importance.

### Stratified LD-score regression

Stratified LD-score regression (sLDSC) was run using the LDSC software^5^ with the baseline v1.2 model, 1000 Genomes Phase 3^68^ European LD scores, HapMap 3 regression SNPs (excluding the HLA region), and 1000 Genomes European allele frequencies. Cell-type-specific annotations were derived from CATlas snATAC-seq binarized peak BED files for the same 111 cell types used by GEMS. A joint model including all 111 cell-type annotations simultaneously alongside the baseline was fit for each trait, so that each cell type’s contribution was assessed after conditioning on all other cell types.

We ranked cell types by the joint-model coefficient *z*-score rather than by the enrichment statistic because the enrichment statistic (proportion of heritability / proportion of SNPs) and its *p*-value reflect a marginal quantity that does not account for shared heritability across correlated cell types, whereas the coefficient captures the unique contribution of each annotation in the joint model. Ranking by the joint-model coefficient *p*-value (Figure S11) or standardized coefficient (*τ ^∗^*, Figure S12) yielded lower silver-standard AUPRCs than the coefficient *z*-score across all four benchmark traits.

### Variant-cell type association

SHAP (SHapley Additive exPlanations) values quantify the contribution of specific cell types to the prediction of GWAS causal status. For each variant, SHAP values were calculated across all cell type tracks, from 100 replicate models and 100 null models for baseline distribution. For each variant-cell type pair, we compared the mean SHAP value obtained from the 100 replicates against the distribution of SHAP values generated from the 100 null models. We calculated a Z-score for every pair and then computed one-tailed p-values to test the hypothesis that the variant’s contribution in a specific cell type was significantly greater than expected by chance. Variant-cell type associations with an FDR-adjusted p-value *<* 0.05 were considered significant.

### Silver-standard enrichment among variant–cell-type associations

To assess whether GEMS variant–cell-type associations are enriched for disease-relevant cell types, we performed one-sided binomial tests for each disease and modality. For a given disease and modality, let *K* denote the total number of cell-type features used by the model (111 for ATAC-seq, 160 for RNA-seq) and *K_s_*the number of those features belonging to the silver-standard set. Under the null hypothesis that variant–cell-type associations are distributed uniformly across cell types, the expected proportion of associations involving a silver-standard cell type is *p*_0_ = *K_s_/K*. We counted the total number of significant variant–cell-type associations *N* (i.e., variant–cell-type pairs with FDR-adjusted *p <* 0.05) and the number of those associations involving a silver-standard cell type *N_s_*. A one-sided binomial test (*H*_1_: observed proportion *> p*_0_) was used to assess whether silver-standard cell types were over-represented among the significant associations. Fold enrichment was computed as (*N_s_/N*) */p*_0_. *P*-values were corrected across all disease–modality combinations using FDR.

### Transcription factor motif disruption analysis

To identify transcription factor (TF) binding motifs disrupted by GWAS variants, we performed input *×* gradient attribution followed by motif discovery for each variant–cell-type pair. For a given variant, Borzoi predicted regulatory output for both reference and alternative allele sequences (524,288 bp centered on the variant) across four independent model folds. Attribution maps were computed using the input *×* gradient method as implemented in gReLU, with predictions aggregated over the cell-type-specific output track(s) and the bins corresponding to the relevant prediction window (16 bins for ATAC-seq at 500 bp; 988 bins for RNA-seq at 31,622 bp). Attribution scores were averaged across the four model folds.

Motif seqlets were discovered independently in the reference and alternative attribution maps within a *±*150 bp window centered on the variant using TF-MoDISco (via the tangermeme implementation). Discovered seqlets were annotated against the HOCOMOCO v12 core motif database, and only seqlets overlapping the variant position were retained. Motifs present in the reference but absent in the alternative attribution were classified as lost by the alternative allele; motifs present only in the alternative were classified as gained. Variants for which TF-MoDISco identified no seqlets (weak attribution signal) were recorded as having no motif change.

### TF family enrichment in MS biclustering modules

To test whether the immune and CNS variant modules identified by biclustering of the MS RNA-seq variant–tissue association matrix (Figure 4B) are driven by distinct TF programs, we assigned each cell in the biclustering heatmap to one of four quadrants defined by the intersection of variant cluster (immune or CNS, from cutting the row dendrogram at *k* = 2) and tissue cluster (immune or CNS, from cutting the column dendrogram at *k* = 2). The full heatmap comprised 3,502 cells (206 variants *×* 17 tissues), of which 1,347 contained at least one TF motif disruption.

TF motifs were mapped to TF families using the HOCOMOCO v12 TFClass annotation. For each TF family and each quadrant, we tested whether cells in that quadrant were enriched for disruptions of that family relative to the remainder of the heatmap using a two-sided Fisher’s exact test, with the 2 *×* 2 contingency table defined as: (cells in quadrant with family disruption, cells outside quadrant with family disruption) *×* (cells in quadrant without, cells outside without). Only families with *≥*3 total disrupted cells were tested.

### cis-Regulatory element enrichment analysis

For each disease, we assembled cell-type-matched cRE datasets from independent published studies: liver ATAC-seq peaks from Currin et al. for LDL; pancreatic islet ATAC-seq peaks from Weng et al. for T2D; CD4^+^ T cell ATAC-seq peaks from Freimer et al. and CD4^+^ T cell caQTL peaks from Mu et al. for ASM; B cell caQTL peaks from Mu et al. for MS memory B cells; and microglia snATAC-seq peaks from Morabito et al. (peaks accessible in *≥*5% of microglia nuclei from a prefrontal cortex AD cohort; 6,895 peaks after liftover to hg19) and microglia H3K27ac ChIP-seq peaks from Li et al. (union set across 15 donors; 99,615 peaks) for MS microglia. All coordinates were mapped to hg19 where necessary.

For each disease, GWAS variants associated with the top GEMS cell type (foreground) were tested for overlap with the matched cRE set relative to the remaining GWAS variants (background) using one-sided Fisher’s exact test. To assess whether GEMS associations extend beyond existing atlas coverage, we repeated each test after removing foreground variants overlapping a CATlas single-cell ATAC-seq peak for the matched cell type.

### Pathway enrichment analysis

To assess whether GEMS RNA-seq variant–cell-type associations place variants near genes in disease-relevant biological pathways, we identified protein-coding genes within the Borzoi RNA prediction window (*±*15,811 bp, corresponding to 31,622 bp total) of each variant using GENCODE annotations. For each disease, variants were split into two groups: a foreground set comprising variants associated with the most frequently associated cell type in the RNA model (ASM: CD4^+^ T cell; LDL: liver; MS: CD14^+^ monocyte; T2D: pancreas), and a background set comprising variants associated with any other cell type but not the foreground cell type. Unique protein-coding genes overlapping the prediction window of each variant set were collected.

For each pathway term from GO Biological Process 2023, we performed a one-sided Fisher’s exact test to assess whether foreground genes were more likely to belong to the pathway than background genes. The 2*×*2 contingency table was constructed as: foreground genes in pathway (*a*), foreground genes not in pathway (*b*), background genes in pathway (*c*), background genes not in pathway (*d*). *P*-values were corrected for multiple testing within each disease–library combination using the Benjamini–Hochberg method.

### Cross-study generalization analysis

To assess whether GEMS and sLDSC cell-type rankings generalize across independent GWAS studies of the same disease, we applied both methods to nine European-ancestry GWAS studies spanning five diseases: asthma (UKB460k), LDL cholesterol (Finucane and Neale UKB studies), multiple sclerosis (IMSGC 2019), type 2 diabetes (Xue 2018, Verma 2024 MVP, Newbury 2025 UKB), and coronary artery disease (van der Harst 2017, Verma 2024 MVP). For each study, joint sLDSC was run on the full GWAS summary statistics using the same 111 CATlas snATAC-seq cell-type peak annotations used in the benchmark analyses. SuSiE fine-mapping was performed separately to identify candidate causal variants (PIP *>* 0.1), which were then processed through the GEMS ATAC-seq pipeline.

For each study, cell types were ranked by the number of GEMS-associated GWAS variants (descending; cell types with zero associations received a tied last rank) and by sLDSC coefficient *z*-scores (descending). Silver-standard AUPRCs were computed by treating the disease-specific silver-standard cell types as positives and all other cell types as negatives, using negative GEMS rank and sLDSC coefficient *z*-scores as scoring functions, respectively (via average_precision_score).

To test whether sLDSC and silver-standard status independently predict GEMS cell-type rankings, we fit a logistic regression model across all nine studies (*N* = 999 cell-type–study observations). The outcome was whether a cell type ranked in the GEMS top 10 (90 positives). The full model was:

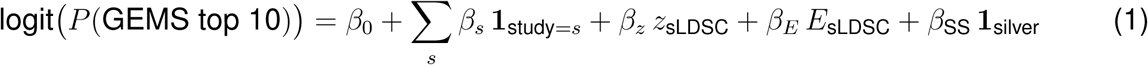

where *z*_sLDSC_ and *E*_sLDSC_ are the sLDSC joint-model coefficient *z*-score and enrichment score (continuous), **1**_silver_ is a binary indicator for silver-standard cell-type status, and the study indicators (dummy-coded, 8 df) absorb disease-level variation because studies are nested within diseases. A likelihood ratio test (LRT) compared the full model to a study-only baseline (3 df). Drop-one LRTs assessed the independent contribution of each predictor conditional on the remaining terms (1 df each).

## DATA ACCESS

Code and data required to replicate this work can be found at Figshare (DOI: 10.6084/m9.figshare.34511559).

## LEAD CONTACT

Requests for further information and resources should be directed to and will be fulfilled by the lead contact, Kipper Fletez-Brant.

## ACKNOWLEDGMENTS

I would like to thank Avantika Lal and Surag Nair for their assistance in troubleshooting and resolving issues with running Borzoi with gReLU. I am also grateful to Will Macnair for sharing a curated list of microglia cis-regulatory element (cRE) datasets, and to Bryce van de Geijn for providing curated GWAS datasets that supported this work.

